# Mu-opioid receptor expression across cell-type specific afferents to the ventral tegmental area in male mice

**DOI:** 10.64898/2026.05.21.726769

**Authors:** Madeline Hohmeister, Oliver Paul Culver, Thomas Jhou

**Affiliations:** Department of Neurobiology, University of Maryland Baltimore, Baltimore, Maryland, United States of America; Department of Neurosciences, Medical University of South Carolina, Charleston, South Carolina, United States of America

## Abstract

The addictive properties of opioids are due in part to these drugs’ ability to alter ventral tegmental area (VTA) activity via activation of mu opioid receptors (MORs) on local and distal inputs. Prior studies have identified numerous opioid-modulated afferents to the VTA, some of which show differing levels of functional modulation by opioids, but the degree to which this parallels differences in receptor expression is not known. Hence, we used retrograde labeling combined with RNAscope to examine oprm1 mRNA expression in VTA-projecting afferents arising from a variety of distal brain regions. Because opioids are thought to be particularly influential on GABAergic afferents to the VTA, we also examined colocalization of oprm1 with GABAergic markers in VTA-projecting neurons. Interestingly, we found that oprm1 mRNA is present in both GABAergic and non-GABAergic VTA-projecting neurons. However, many (though not all) GABAergic afferents expressed higher levels of oprm1 compared to most non-GABAergic afferents (especially those arising from the cortex). These results complement previous anatomical studies that had examined oprm1 expression in these regions but in a non-quantitative way and without regard to their efferent targets. Our findings encourage future work to examine the functional implications of MOR sensitivity within these afferent pathways.

## Introduction

Agonists of the mu opioid receptor (MOR) produce strong rewarding effects that can lead to development of opioid use disorder (OUD), which has become increasingly prevalent in recent years [1,2,3]. MOR agonists are widely recognized to influence reward behavior by inhibiting GABAergic inputs to midbrain dopamine (DA) neurons in the ventral tegmental area (VTA), thereby disinhibiting DA release. Furthermore, early studies suggested that GABAergic neurons within the VTA itself are the primary target of MOR agonists [4,5]. However, subsequent studies found that MOR agonists exert stronger influences on GABAergic afferents arising from regions outside the VTA, such as the rostromedial tegmental nucleus (RMTg) and nucleus accumbens (NAc) [6]. Functional evidence for MOR-induced inhibition of afferents arising from the NAc, RMTg, and VTA itself have been supported by anatomical evidence confirming the presence of mRNA for *oprm1* (the gene encoding MORs) in these brain regions [7,8].

Notably, our knowledge of opioiderigc influences on different GABAergic VTA afferents is incomplete. For example, functional studies show that MOR agonists more strongly inhibit GABAergic currents in VTA DA neurons due to stimulation of afferents arising from the RMTg than from other GABAergic sources such as NAc or VTA itself [6]. However, it is unknown whether these differences might be related to differences in MOR expression levels. While anatomical studies subsequently identified high levels of MOR-expression in RMTg, they did not address whether this high expression occurs specifically in neurons projecting to the VTA, versus other targets of RMTg efferents [7]. Furthermore, additional sources of GABAergic input to the VTA have been reported [9] and we now know that many of these regions are also influenced by MOR agonists. For example, recent studies have shown that MOR agonists also inhibit GABAergic inputs to VTA neurons arising from the VP [10]and BNST [11]. However, MOR expression within these and other distal GABAergic VTA afferents is not as well studied. Gaps in our current knowledge of how opioids modulate inhibitory input to the VTA could be addressed by a comprehensive study broadly examining which VTA-projecting GABA neurons arising from which brain regions express MORs and to what degree.

It is important to note that VTA neurons can be modulated by multiple afferent pathways arising from different brain regions and expressing different neurotransmitters [9]. While many studies have focused on how MOR agonists affect GABAergic VTA inputs due to the VTA-DA disinhibition hypothesis, the functional role of MORs on non-GABAergic inputs should not be overlooked. Additional functional and anatomical studies also show that MORs target non-GABAergic afferents, such as those expressing glutamate, to promote reward behaviors [12,13,14]However, other studies have shown that application of MOR agonists do not alter excitatory glutamate release onto VTA DA neurons [15], suggesting that non-GABAergic inputs lack traditional Gi-coupled MORs. Hence, more anatomical evidence regarding the extent of MOR expression among non-GABAergic afferent pathways (especially compared to MOR expression among GABAergic afferent pathways) is needed.

The present study aims to quantify MOR expression across a broad range of GABAergic and non-GABAergic pathways that specifically project to the VTA. Utilizing retrograde tracers combined with RNAscope fluorescent in situ hybridization, we found that *oprm1* expression is widespread throughout multiple GABAergic afferents to the VTA, with some significant differences between specific afferents such as accumbens, hypothalamus, and RMTg. Oprm1 is also present on non-GABAergic afferents, albeit to a lesser degree in most cases. These findings bolster previous functional and anatomical studies investigating the effect of MOR agonists on VTA function and invite future research into the clinical implications of MOR functions within these afferent pathways.

## Methods

### Animals

Adult male C57BL/6J mice were obtained from Jackson Laboratory (cat# 000664) and housed in standard ventilated cages within a temperature-controlled (21℃), 12/12 h light/dark cycle (lights on at 6:00 am) vivarium, with food and water provided ad libitum. Procedures conformed to the National Institute of Health Guide for the Care and Use of Laboratory Animals, and all protocols were approved by the University of Maryland Baltimore Institutional Animal Care and Use Committee.

### Stereotaxic surgery

Mice were anesthetized via inhaled isoflurane (5% induction, 1–3% maintenance), placed into a stereotaxic apparatus, and administered the analgesic carprofen (5 mg/kg). Intracranial injections were made using a pulled glass micropipette and Nanoliter Microinjection Pump (R-480, RWD Life Science); injections were allowed to diffuse for 5 min prior to retracting glass pipet injector. The retrograde tracer (Red Retrobeads^TM^, Lumafluor Inc.) was injected unilaterally in the VTA of mice (250–500 nl, AP −3.1, ML +1.3, DV −4.8 from dura, 10° angle).

### Sacrifice and Tissue Collection

3-7 days after surgery, mice were anesthetized via inhaled isoflurane and rapidly decapitated. The brains were immediately frozen in isopentane and cut into 40-μm sections on a cryostat. For each brain region of interest, 3 serial sections were selected and mounted on glass slides. All tissue was stored at -80℃ until RNAscope in situ hybridization assays were run.

#### RNAscope Fluorescent in Situ Hybridization

Fluorescent in situ hybridization assays were performed using the RNAscope Multiplex Fluorescent Reagent Kit v2 (Advanced Cell Diagnostics, ref# 323110) according to the manufacturer’s instructions. Briefly, tissue sections from each brain region of interest were fixed in a 10% formalin solution for 15 minutes, then dehydrated with ethanol and pretreated with a 1:4 diluted solution of protease III (Advanced Cell Diagnostics, ref# 322337). Tissue sections that included the RMTg were hybridized with RNAscope probes specific for foxp1 (Advanced Cell Diagnostics, ref# 485221-C2) and oprm1 (Advanced Cell Diagnostics, ref# 315841-C3), while all other sections were hybridized with probes specific for gad2 (Advanced Cell Diagnostics, ref# 439371-C2) and oprm1(Advanced Cell Diagnostics, ref# 315841-C3). Hybridization was amplified for detection using the TSA fluorophores for fluorescein (498) (Akoya Biosciences, ref# TS-000200) and far red (647) (Akoya Biosciences, ref# NEL705A001KT) and stained with DAPI (Advanced Cell Diagnostics, ref# 323108) to label the nucleus of cells. A negative control RNAscope assay, in which hybridization of the oprm1 probe was excluded from the protocol, was also run on all relevant brain sections in order to control for non-specific binding and false signals in our oprm1 channel (Supplemental Fig. 1).

**Supplemental Fig 1.**
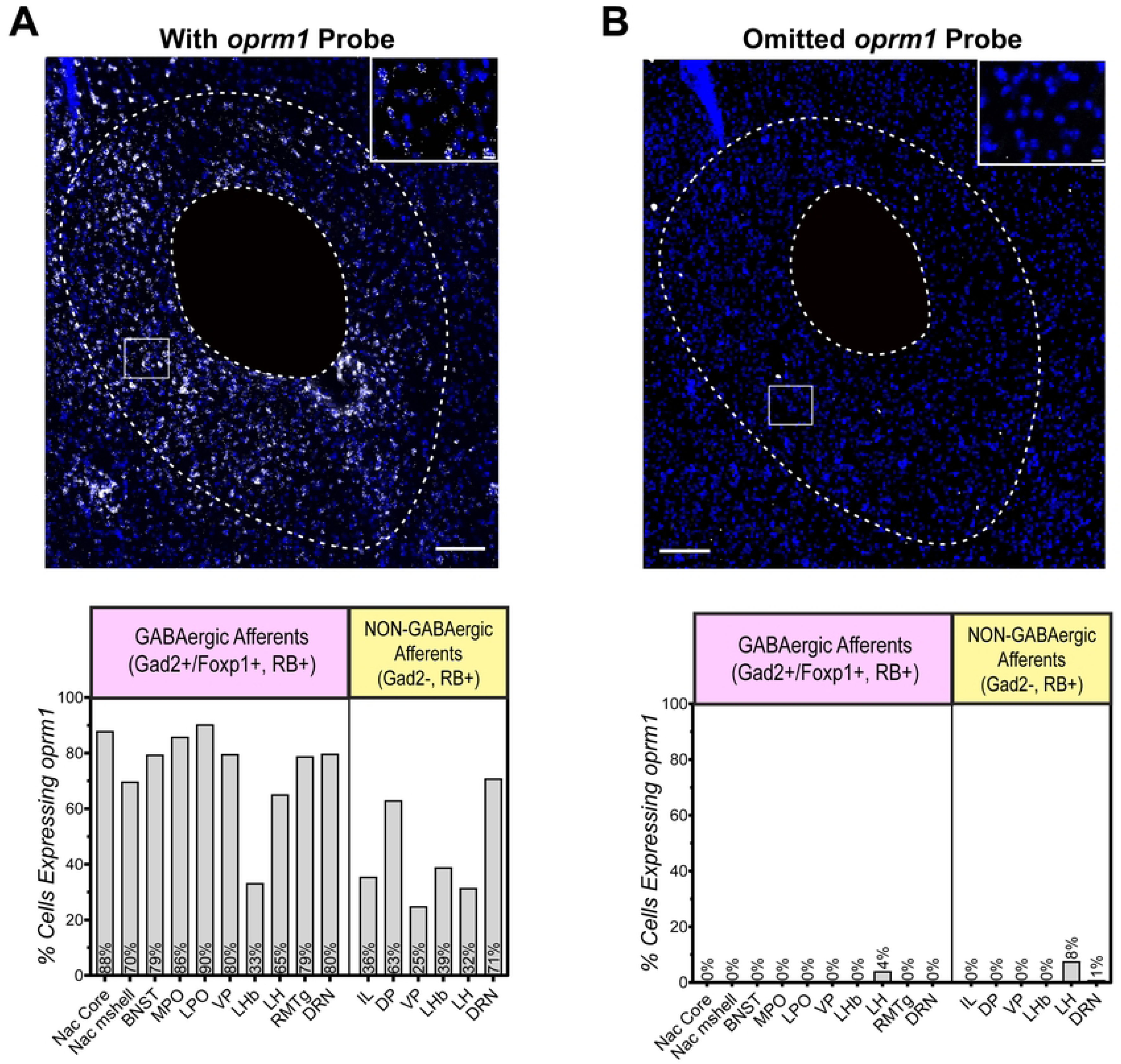
*oprm1* probe negative control to assess non-specific binding A negative control RNAscope assay, in which the *oprm1* probe was excluded from the in-situ hybridization protocol, was run on all 12 regions of interest from animal #1 to control for non-specific signals. (A) 20x representative image showing RNAscope signal in the NAc core when the *oprm1* probe was included, versus when it was omitted (B). (C) Quantification of percent of GABAergic or non-GABAergic retrogradely labeled cells that co-express *oprm1* in each brain region of interest for Animal #1 when the *oprm1* probe was included, versus omitted (D). Scale bars = 100µm for 20x image, and 10µm for magnified inset image.

### Image Acquisition and Analysis

For each brain region that we analyzed, three serial tissue sections were imaged on a MICA Microhub (Leica Microsystems), using a 20x objective. Image processing and colocalization analysis were carried out using the open-source package Fiji, which is built on ImageJ software developed by the National Institutes of Health. Images of each fluorescent channel (gad2/foxp1 channel, oprm1 channel, and RetroBead channel), were first processed with a spatial bandpass filter created by subtracting the image processed with a gaussian blur of sigma (radius) 2 from the same image processed with a gaussian blur of sigma (radius) 1. This preprocessing was followed by thresholding, binary image conversion, and water-shedding via ImageJ’s classic watershed plugin in order to eliminate background noise and isolate fluorescent mRNA puncta labeling.

For colocalization analysis, brain regions of interest were first delineated in reference to existing mouse brain atlases (Paxinos and Franklin, 2001 and Alan Institute for Brain Science, 2004). Brain regions of interest were included if we observed strong retrograde labeling in at least 3 of the mice assayed. Within each brain region, we defined regions of interest (ROIs) as elliptical regions encompassing individual retrogradely labeled (RB +) cells, and the particle function in ImageJ was used to quantify the number of gad2 (or foxp1 for the RMTg) and oprm1 puncta within each cell/ROI. These values were exported to an excel spreadsheet that was further processed using an R scripts, in order to categorize retrogradely labeled cells as either GABAergic and oprm1 positive (gad2+, oprm1+, RB+), GABAergic and oprm1 negative (gad2+, oprm1-, RB+), non-GABAergic and oprm1 positive (gad2-, oprm1+, RB+), or non-GABAergic and oprm1 negative (gad2+, oprm1+, RB+). For RMTg sections, retrogradely labeled cells were categorized as either GABAergic and oprm1 positive (foxp1+, oprm1+, RB+), GABAergic and oprm1 negative (foxp1+, oprm1-, RB+), non-GABAergic and oprm1 positive (foxp1-, oprm1+, RB+), or non-GABAergic and oprm1 negative (foxp1+, oprm1+, RB+) Cells were considered GABAergic if they expressed >3 gad2 or foxp1 puncta, and oprm1 positive if they expressed >5 oprm1 puncta (Fig 2.).

### Statistical Analysis of oprm1 Puncta Per Cell

Statistical analysis of oprm1 transcript levels was performed using Prism v9.0.0 (GraphPad Software LLC). Oprm1 puncta per cell counts for each afferent pathway were normalized using a square root transform. Normality was assessed using the D’Agostino and Pearson test. The Welch ANOVA test and Games-Howell (alpha=0.05) post hoc test were used to compare the average number of oprm1 puncta per cell between different afferent pathways in individual animals The Welch ANOVA test and Dunnett’s T3 (alpha=0.05) post hoc test were used to compare the average number of oprm1 puncta per cell between different afferent pathways at a group level.

## Results

### Anatomical Organization of Major Afferents to the VTA

VTA-projecting cells from across the brain were retrogradely labeled with red RetroBead fluorescent latex microspheres injected into the VTA of C57BL/6 mice (Fig 1A). Microspheres were chosen over other fluorescent retrograde tracers based on pilot studies showing that the fluorescent signal from the microspheres was not degraded by subsequent RNAscope *in situ* hybridization histochemical procedures. We placed injections into 14 animals, of which 9 were rejected due to substantial tracer spread outside of the VTA. This left five mice with injections confined to the VTA, and with limited spread to adjacent regions such as the substantia nigra par reticulata (SNr) or interpeduncular nucleus (IPN) (Fig 1B). Three to seven days after injection, animals were sacrificed, and brains processed histologically. Retrogradely labeled cells were then examined throughout the brain [9,16] and quantified using boundaries prescribed by a standard mouse brain atlas [17] (Fig 1 C-D).

**Fig 1.**
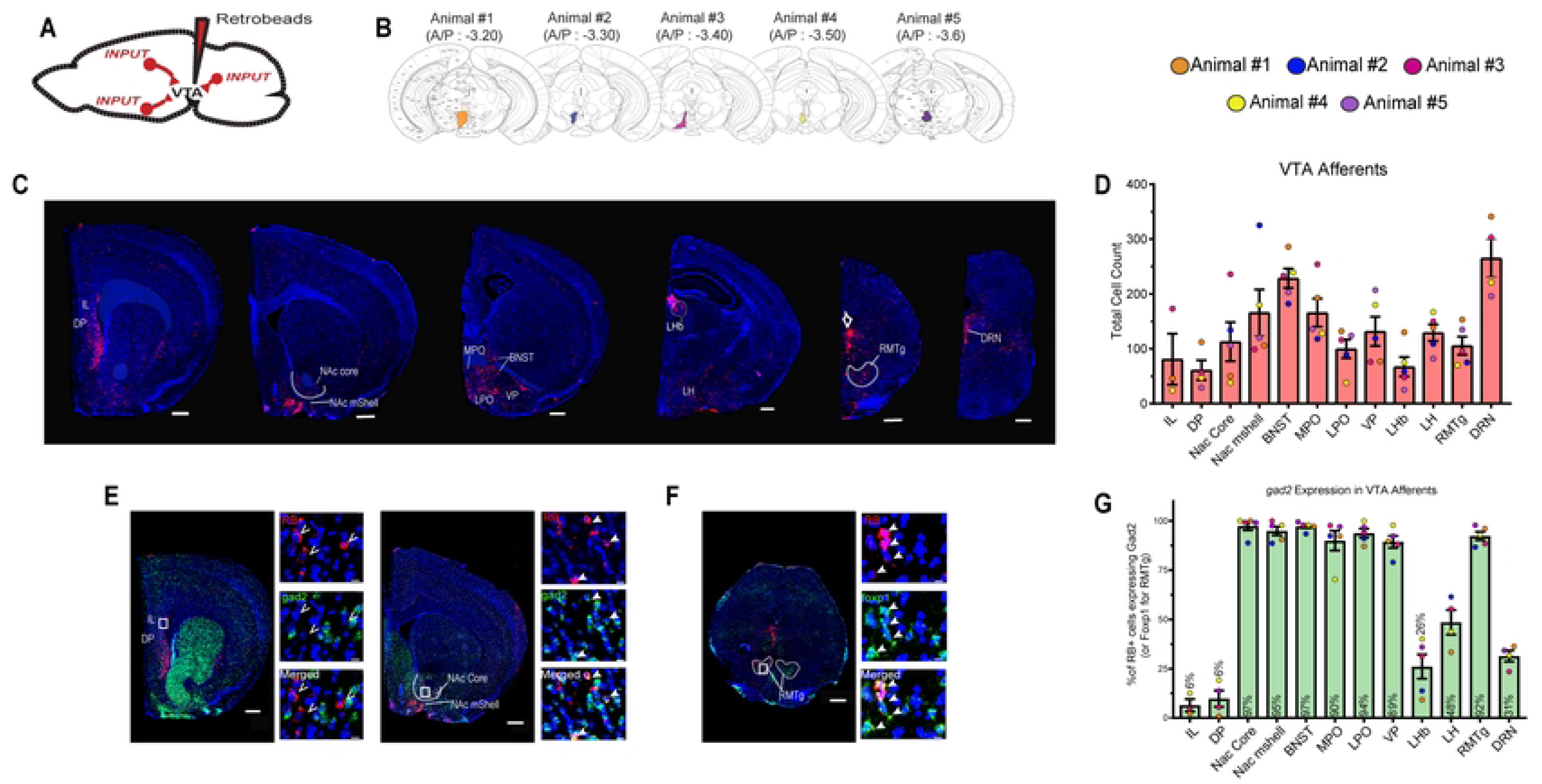
Retrograde Mapping of Cell-Type Specific Afferents to the VTA (A) Schematic of RetroBead (RB) fluorescent latex microsphere injection into the VTA. (B) RB distribution in VTA injection sites of 5 C57BL/6 male mice. (C) 20x representative images (from animal #3) showing RB (red) labeling to different brain regions of interest, alongside DAPI stain (blue) to show cytoarchitecture. Scale bar = 500µm. (D) Quantification of RB labeled cells in each brain region of interest (E) 20x representative images showing RB retrograde label (red) alongside RNAscope signal for *gad2* mRNA (green) in accumbens and cortex. Scale bar = 500µm. High-magnification insets show RB-labeled cells co-expressing *gad2* (filled arrows) or lacking *gad2* (unfilled arrows). Scale bar = 10µm. (F) Same as D, but in RMTg brain region, and the green label indicates *foxP1,* which overlaps strongly with *gad2* in RMTg, and delineates the RMTg outline more clearly than *gad2*. (G) Mean ± SEM of the percent of GABAergic afferents (*gad2*/*foxp1*+, RB+ cells) in each brain region relative to total RB+ cells for that region. Each data point represents the percentage for one animal. Abbreviations of brain regions: IL: infralimbic cortex (n=3), DP: dorsal peduncular cortex (n=4 mice), NAc core: nucleus accumbens core (n=5 mice), NAc mshell: nucleus accumbens medial shell (n=5 mice), BNST: bed nucleus of the stria terminalis (n=5 mice), MPO: medial preoptic area (n=5 mice), LPO: lateral preoptic area (n=5 mice), VP: ventral pallidum (n=5 mice), LHb: lateral habenula (n=5 mice), LH: lateral hypothalamus (n=4 mice), RMTg: rostromedial tegmental nucleus (n=5 mice), DRN: dorsal raphe nucleus (n=4 mice).

We identified 12 brain regions that consistently contained neurons with strong retrograde labeling. These were: the infralimbic cortex (IL), dorsal peduncular cortex (DP) nucleus accumbens core and medial shell (NAc core and NAc mshell respectively), bed nucleus of the stria terminalis (BNST), medial preoptic and lateral preoptic area (MPO and LPO respectively), ventral pallidum (VP), lateral habenula (LHb), lateral hypothalamus (LH), rostromedial tegmental nucleus (RMTg), and dorsal raphe nucleus (DRN) (Fig 1C-D). The presence of strong labeling in these structures is largely consistent with previous anatomical and functional studies [9,16].

### Cell-Type Specific Anatomical Organization of Major Afferents to the VTA

We used retrograde tracing combined with RNAscope *in situ* hybridization for the *gad2* gene (which encodes one of two major GABA-synthesizing enzymes in the brain) to determine what proportion of VTA-projecting neurons in each brain region were GABAergic (Fig 1E-G). *Gad2* was chosen as our GABAergic marker over its isoform *gad1* because of *gad2*’s preferential role in the synthesis of GABA within synaptic terminals where they would contribute to vesicular release [18,19,20].

Only a small proportion of retrogradely labeled cells arising from the IL, DP, LHb and DRN expressed *gad2* mRNA, aligning with previous findings that the IL, DP, and LHb predominantly send glutamatergic projections to the VTA [9, 13, 21] while the DRN predominantly send glutamatergic and serotonergic projections [22]. In contrast, a majority of VTA-projecting cells arising from the NAc core, NAc mshell, BNST, MPO, LPO, VP, and LH, co-expressed *gad2* mRNA (Fig. 1E, G).

We used *foxp1* instead of *gad2* to identify GABAergic projection neurons arising from the RMTg, because it is expressed at higher levels in the RMTg than its surroundings (Fig 1F), unlike *gad2* which is abundant throughout. Previous studies have shown that a majority of GABA cells in the RMTg (a predominantly GABAergic structure itself) co-express *foxp1* mRNA [23]. Consistent with these studies, we found that a majority of retrogradely labeled cells within the RMTg expressed *foxp1* (Fig. 1G).

### Expression of oprm1 Across Major GABAergic and Non-GABAergic Inputs to the VTA

Our findings are consistent with previous findings that the VTA receives input from multiple distal brain regions [9,16]. However, the anatomic distribution of *oprm1* expression across many of the input pathways we have identified is unknown. To answer this question, we again used retrograde tracing combined with RNAscope to examine expression of *oprm1* in VTA-projecting GABAergic and non-GABAergic neurons across all brain regions of interest.

Similar to previous studies, a large proportion of VTA-projecting GABA cells arising from the NAc [8] and RMTg [7] co-expressed *oprm1* (Fig 2 A-B). We found similarly high proportions of *oprm1* in VTA-projecting GABA neurons arising from other distal inputs. In fact, *oprm1* mRNA was distributed across every inhibitory projection pathway we investigated, including the subregions of the NAc, BNST, MPO, LPO, VP, LH, and DRN (Fig 2A-B).

**Fig 2.**
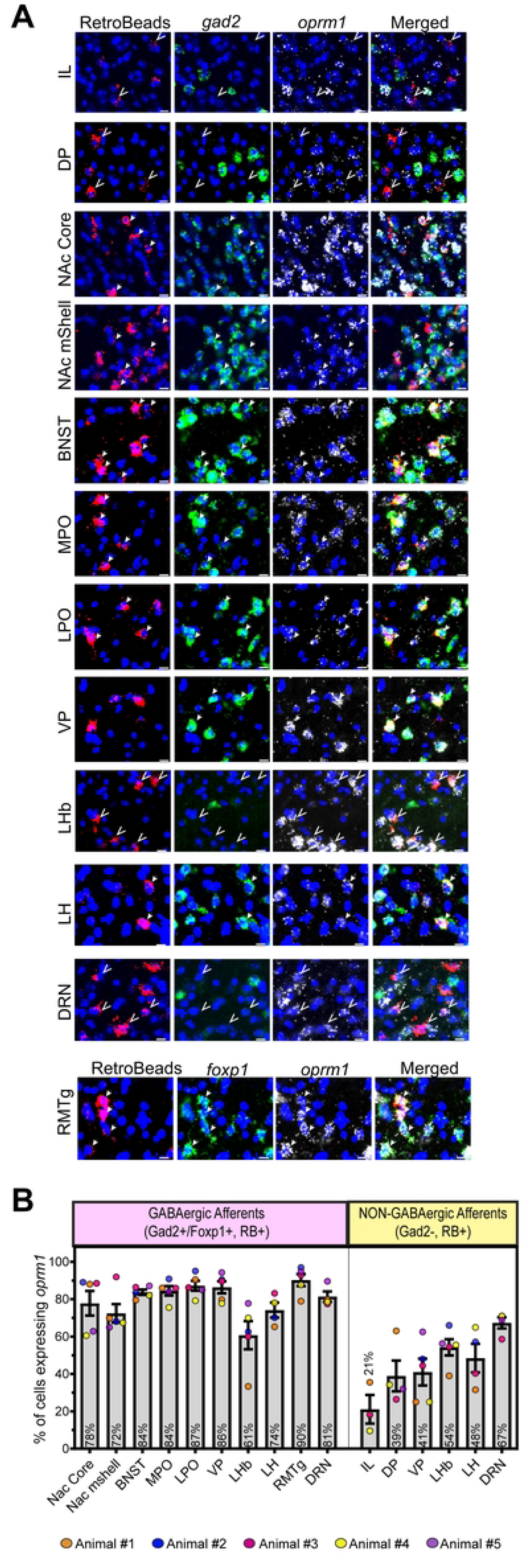
*opmr1* Expression Across Cell-Type Specific Afferents to the VTA (A) High-magnification representative images of all 12 brain regions of interest. Columns 1-4 indicate: RB retrograde label (red), *gad2* mRNA (green), *oprm1* mRNA (white), and merged images. In the bottom row (RMTg images), green signal indicates *foxP1* instead of *gad2*. DAPI counterstain is shown in blue in all images. Filled arrows indicate neurons co-expressing all three of the markers RB, *gad2/foxp1*, and *oprm1*, i.e. while unfilled arrows indicate cells expressing RB and *oprm1*, but not *gad2*/*foxp1*, i.e. non-GABAergic VTA afferents. (B) Percent (mean ± SEM) of GABAergic or non-GABAergic retrogradely labeled cells that co-express *oprm1* in each brain region of interest. Each point represents one animal. Percent *oprm1* expression in GABAergic populations = percent of *gad2*+/*foxp1*+, RB+ cells that co-express *oprm1*. Percent *oprm1* expression in non-GABAergic populations = percent of *gad2*-/*foxp1*-, RB+ cells that co-express *oprm1*. Scale bar = 10µm. Group sizes: IL (n=3 mice), DP (n=4 mice), NAc core (n=5 mice), NAc mshell (n=5 mice), BNST (n=5 mice), MPO (n=5 mice), LPO (n=5 mice), VP (n=5 mice), LHb (n=5 mice), LH (n=4 mice), RMTg (n=5 mice), DRN (n=4 mice).

To assess whether MORs are also expressed on non-GABAergic projections to the VTA, we looked at *oprm1* expression in regions where >10% of retrogradely labeled cells lacked *gad2* in a majority of animals. Expression of *oprm1* puncta was observed in a moderate proportion of VTA-projecting non-GABAergic neurons arising from the IL, DP, VP, LHb, LH and DRN (Fig 2 A-B.).

After observing high frequencies of *oprm1* colocalization across multiple input pathways, we examined *oprm1* mRNA transcript levels by quantifying the number of fluorescent puncta per cell (Fig 3). We then compared the average number of *oprm1* puncta per cell between different afferent populations. Due to the possibility that variations in bead placement within the VTA may have resulted in retrograde labeling of different afferent populations between animals [9], we first analyzed animals individually (Fig 3B-F).

**Fig 3.**
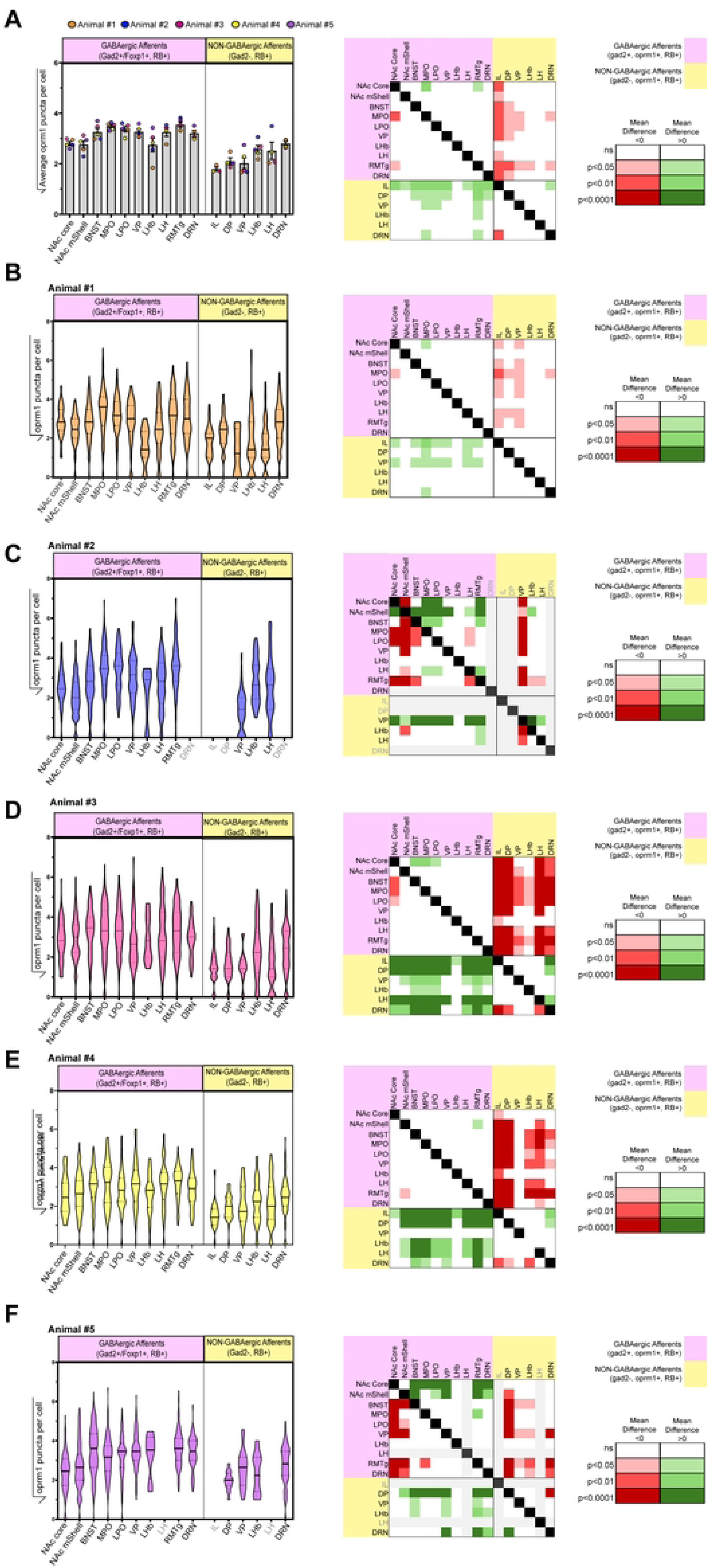
Differences in oprm1 Transcript levels Across Different VTA Afferent Populations (A-F) *oprm1* puncta per cell across GABAergic (pink) and NON-GABAergic (yellow) afferents to the VTA arising from different brain regions at a group level (A), where each point represents the average puncta per cell count for an individual animal, and across individual animals (B-F), where violin plots represent distribution of puncta counts for individual cells after square root transform to normalize data. Heatmaps show statistically significant differences in the average number of puncta per cell between different afferent populations, using Welch ANOVA test followed by Dunnett’s T3 multiple comparisons for group level analysis and Games-Howell multiple comparisons for individual animals (alpha 0.05). Shading from light to dark indicates increasing significance (p<0.05, p<0.01, p<0.0001), while color indicates direction of difference, with red indicating column population has less *oprm1* than row population, while green indicates column population expresses greater *oprm1* than row population, and white indicates no difference. Gray columns indicate that afferents arising from that brain region were unable to be quantified for that animal. Group level Welch ANOVA; W=20.15, p<0.0001, IL=3 mice, DP=4 mice, NAc core=5 mice, NAc mshell=5 mice, BNST=5 mice, MPO=5 mice, LPO=5 mice, VP=5 mice, LHb=5 mice, LH=4 mice, RMTg=5 mice, DRN=4 mice. Animal #1 Welch ANOVA; W=24.51, p<0.0001, IL=45 cells, DP=111 cells, NAc core=50 cells, NAc mshell=95 cells, BNST=278 cells, MPO=178 cells, LPO=115 cells, VP=69 (GABA),8 (Non-GABA) cells, LHb=13 (GABA), 125 (Non-GABA) cells, LH=46 (GABA), 92 (Non-GABA) cells, RMTg=147 cells, DRN=124 (GABA), 217 (Non-GABA) cells. Animal #2 Welch ANOVA; W=25.37, p<0.0001, NAc core=119 cell, NAc mshell=288 cells, BNST=166 cells, MPO=109 cells, LPO=81 cells, VP=94 (GABA), 35 (Non-GABA) cells, LHb=8 (GABA), 50 (Non-GABA) cells, LH=67 (GABA), 35 (Non-GABA) cells, RMTg=65 cells. Animal #3 Welch ANOVA; W=43.67, p<0.0001, IL=165 cells, DP=53 cells, NAc core=234 cells, NAc mshell=99 cells, BNST=228 cells, MPO=248 cells, LPO=109 cells, VP=76 (GABA), 9 (Non-GABA) cells, LHb=15 (GABA), 33 (Non-GABA) cells, LH=83 (GABA), 67 (Non-GABA) cells, RMTg=84 cells, DRN=71 (GABA), 232 (Non-GABA) cells. Animal #4 Welch ANOVA; W=16.95, p<0.0001, IL=21 cells, DP=38 cells, NAc core=38 cells, NAc mshell=180 cells, BNST=190 cells, MPO=90 cells, LPO=149 cells, VP=63 (GABA), 9 (Non-GABA) cells, LHb= 30 (GABA), 45 (Non-GABA) cells, LH=73 (GABA), 94 (Non-GABA) cells, RMTg=65 cells, DRN=70 (GABA), 151 (Non-GABA) cells. Animal #5; W=28.03, p<0.0001, DP=25 cells, NAc core=105 cells, NAc mshell=117 cells, BNST=202 cells, MPO=127 cells, LPO=115 cells, VP=188 (GABA), 19 (Non-GABA) cells, LHb=8 (GABA), 17 (Non-GABA) cells, RMTg=132 cells, DRN=67 (GABA), 129 (Non-GABA) cells.

Among GABAergic inputs to the VTA, afferents from the NAc core had fewer *oprm1* puncta per cell compared to GABAergic afferents arising from the MPO in four out of five animals (Fig 3B, C, D and F). Additionally, afferents from the NAc mshell had fewer *oprm1* puncta per cell compared to GABAergic afferents arising from the RMTg in three out of five animals (Fig 3C, E and F). Afferents arising from the different regions of the NAc also had significantly fewer oprm1 puncta per cell compared to GABAergic afferents arising from the BNST and LPO in 3 out of five animals (Fig 3C, D and F).

Compared to a majority of GABAergic afferents, non-GABAergic cortical afferents arising from the IL and DP had significantly fewer *oprm1* puncta per projection neuron. This difference was robust and present in all animals where tissue for these regions was available (Fig 3B, D, E, and F). Non-GABAergic projection neurons arising from the VP, LHb, LH and DRN also displayed significantly fewer *oprm1* puncta per cell compared to multiple GABAergic afferents in a majority of animals. However, which GABAergic afferents each of these non-GABAergic afferents differed from and to what degree varied between animals (Fig 3B-F).

When comparing among GABAergic regions at a group level (n=5 mice), we saw lower *oprm1* levels in GABAergic projection neurons arising from the NAc core compared to those arising from the MPO (p=0.0058, n=5 mice) and RMTg (p=0.028, n=5 mice),while differences in *oprm1* puncta per cell did not reach significance for other pairs of GABAergic afferent populations. (Fig. 3A).

When comparing GABAergic to non-GABAergic afferents at a group level, we saw significantly lower *oprm1* levels among non-GABAergic projection neurons arising from the IL compared to GABAergic projection neurons arising from the NAc core (p=0.0071), NAc mshell (p=0.0408, n=5 mice), and LH (p=0.0294) We also saw significantly lower *oprm1* levels among non-GABAergic projection neurons arising from both the IL and DP compared to GABAergic projection neurons arising from the BNST (IL: p=0.0041, DP: p=0.0254), MPO (IL: p=0.0017, DP: p=0.0158), LPO (IL: p=0.0008, DP: p=0.011), VP (IL: p=0.001, DP: p=0.0162), RMTg (IL: p=0.0005, DP: p=0.0057), and DRN (IL: p=0.0056, DP: p=0.0305). Additionally, group level analysis showed that there were significantly lower *oprm1* levels among non-GABAergic VTA projecting neurons arising from both the VP and DRN compared to GABAergic projection neurons arising from the MPO (VP: p=0.0355, DRN: p=0.0203) and RMTg (VP: p=0.0245, DRN: 0.0333). Non-GABAergic LHb projection neurons also had significantly lower *oprm1* levels compared to GABAergic RMTg projection neurons (p=0.0247). These non-GABAergic LHb afferents also appeared to have lower *oprm1* levels compared to GABAergic MPO afferents, though this difference did not quite reach significance (p=0.0627). Finally, GABAergic afferents arising from the LH did not significantly differ from any other afferents we surveyed; however, this may have been due to high variability in average *oprm1* levels among different animals within this region (Fig 3A).

## Discussion

In this study, we used retrograde labeling combined with RNAscope fluorescent in situ hybridization to examine *oprm1* mRNA expression in VTA-projecting neurons arising from multiple brain regions. We established that MORs are present in GABAergic VTA projection neurons arising from multiple distal brain regions outside of the VTA including the NAc, RMTg, VP, BNST, LPO, MPO, LH, and DRN. This finding supports previous findings that MOR agonists target GABAergic neurons projecting to the VTA from a variety of brain regions outside of the VTA itself [6, 10,11, 24, 25]

At both a group level and within a majority of individual animals, *oprm1* transcript levels differed significantly among GABAergic afferents arising from the NAc compared to those arising from both the MPO and RMTg. This finding is consistent with previous functional ones showing that some inputs to the VTA may be more affected than others by the action of opioids [6]. When comparing afferent populations at an individual level, several differences, e.g. the higher level of expression in RMTg afferents relative to accumbens, were seen in only 3 out of 5 animals (with the remaining two trending in the same direction, but without showing significance), possibly due to variations in tracer placement within the VTA. Locations of our VTA tracer injections differed in both anterior-posterior and mediolateral placement, and these differences have been shown to alter which afferent populations are retrogradely labeled [9]. This subsequently caused some animals to have lower numbers of retrogradely cells in certain brain regions, reducing statistical power for comparisons made in those animals as well.

We also identified *oprm1* expression in non-GABAergic VTA-projection neurons arising from brain regions previously reported to make excitatory connections with the VTA, including IL, DP, VP, LHb, LH and DRN. Our findings further uncover non-GABAergic inputs outside of the VTA that also express *oprm1*. Of note, we did observe significantly lower levels of oprm1 transcripts in non-GABAergic afferents (especially cortical ones) compared to GABAergic afferents. This finding provides a possible explanation as to why some studies have claimed that MOR agonists have little to no effect on some glutamatergic inputs to the VTA [15]. However, the presence of oprm1 mRNA transcript expression in non-GABAergic afferents nonetheless invites future investigation into how MOR agonists may alter these important inputs to the VTA.

Because our RNAscope experiments only probed for *gad2* and *oprm1* mRNA, we were unable to determine whether non-GABAergic projection cells expressed glutamatergic markers, versus other neurotransmitter markers, an important consideration for understanding how they impact VTA activity. While the LHb is considered to be largely glutamatergic [21, 26], other sites such as the DRN likely send both glutamatergic and serotonergic (in addition to GABAergic) projections to VTA dopamine neurons [22]. Further functional studies are also needed to determine if MOR agonists impair the ability of these non-GABAergic pathways to regulate VTA activity and subsequent reward learning.

While our study identifies distinct *oprm1* expressing VTA-projecting pathways, we are unable to address whether these neurons innervate DA or non-DA neurons in the VTA. Indeed, previous studies have found that many of these distal inputs can innervate both VTA GABA and dopamine neuron subpopulations [9]. Hence, MORs could disinhibit multiple types of VTA neurons, and future studies will need to determine which ones are influenced by MOR-mediated modulation of the GABAergic inputs we identified.

While we did not examine behavioral effects of opioids in the brain regions we examined, prior studies showed that many of them could drive motivated behaviors. For example, injection of morphine into the LH stimulates food intake [27], while injection of the MOR antagonist naloxone into the MPO inhibits CPP produced by sexual reinforcement [28]. However, additional behavioral studies will be needed to test whether these effects are due specifically to projections from these regions to the VTA.

One final shortcoming of this study is that we only examined MOR expression in male mice, whereas previously studies reported differences in MOR expression in males compared to cycling females [29,30,31]. Future studies will be needed to address whether males and females express differing levels of oprm1 mRNA among the distal VTA afferents we identified in this study.

